# Apical-basal polarity regulates Collagen IV-dependent cell-cell adhesion in the *Drosophila* adipose tissue

**DOI:** 10.1101/2025.04.17.649307

**Authors:** Jameela Almasoud, Cyril Andrieu, Bren Hunyi Lee, Anna Franz

## Abstract

E-Cadherin-based adherens junctions are the main type of junctional compartment in epithelia mediating cell-cell adhesion. The apical-basal cell polarity machinery positions adherens junctions at the apical-lateral border. The *Drosophila* adipose tissue, called the fat body, forms a monolayer in which cells adhere to each other through the binding of Integrin to pericellular Collagen IV concentrations. It is currently not known, how these atypical adhesion complexes form. Here we examine the epithelial apical-basal cell polarity network in the larval fat body. We find that this tissue displays an apical-basal cell polarity, with the apical proteins aPKC, Crumbs and Par-6 and the basolateral proteins Lgl and Dlg on opposite sides. Crumbs, aPKC, Scribble, and Lgl knockdown in the fat body leads to cell-cell-adhesion defects. We find that aPKC plays a key role in mediating cell-cell adhesion by regulating the formation of Collagen IV-Intercellular-Adhesion-Concentrations. We further show that during fat body remodeling, the steroid hormone Ecdysone regulates the loss of apical-basal polarity and Collagen IV-Intercellular-Adhesion-Concentrations to induce cell-cell dissociation and initiation of amoeboid swimming cell migration. Our work hence uncovers a novel role for the apical-basal polarity machinery in the *Drosophila* adipose tissue in regulating cell-cell adhesion via Collagen IV-Intercellular-Adhesion-Concentrations.

## Introduction

Mesoderm-derived adipocytes form true tissues by tightly associating with each other. In mammals the adipose tissue is distributed over multiple subcutaneous and visceral depots [1]. In *Drosophila* larvae, adipocytes are large polyploid cells that are organized in a single continuous monolayer, called the fat body. This tissue lies inside the body cavity and is surrounded by hemolymph, the body fluid. The fat body tissue bifurcates at the anterior end of the animal into two sheets that extend on each side towards the posterior end of the animal surrounding the internal organs including the gut like a bilateral apron. The mechanisms that maintain adipose tissue architecture and its functional significance remain largely elusive. In contrast, the tissue architecture of epithelia and its function have been extensively studied. In *Drosophila*, cell-cell adhesion through E-Cadherin-based adherens junctions at the apical-lateral border mediates the formation of sheets which is dictated by apical-basal cell polarity [2]. A conserved set of polarity proteins determines the various domains in epithelial cells as uncovered mostly through studies in *Drosophila* and *C. elegans*. These showed that the apical domain is specified by the transmembrane protein Crumbs, the adaptor protein Stardust, and the Par-6/atypical protein kinase C (aPKC) complex. Bazooka (Baz in flies, named Par-3 in other organisms) defines the boundary between the apical and lateral domains. It plays a key role in positioning the apical adherens junctions and localising the apical factors, which then exclude Baz from the apical domain [3–8]. Discs large (Dlg), Lethal (2) giant larvae (Lgl) and Scribble (Scrib) mark the rest of the lateral domain and the basal domain [9]. Mutual antagonism between apical and lateral factors then ensures the maintenance of the identity of the apical and lateral domains [10, 11]. Moreover, a basement membrane (BM) composed of the extracellular matrix proteins Collagen IV, Perlecan, Nidogen and Laminin underlies the basal domain of epithelial cells [12].

In contrast to epithelia, the fat body is not known to have an apical-basal cell polarity. Cell-cell adhesion here has been proposed to be mediated by two alternative mechanisms, via E-Cadherin-based adherens junctions [13] or via Collagen IV-Intercellular-Adhesion-Concentrations [14]. In agreement with a role of E-Cadherin in fat body cell-cell adhesion, it was reported that E-Cadherin is localized at cell-cell vertices in the larval fat body, and this localization is then lost during fat body remodeling as cells dissociate [13]. More recently, it was shown that extracellular Collagen IV-containing punctae are found spread along the cell-cell vertices in the pericellular space of the larval fat body [14]. Neighboring cells attach to these Collagen IV-Intercellular-Adhesion-Concentrations (CIVICs) via Integrin and Syndecan receptors which is essential for cell-cell adhesion [14]. However, it remains unknown how CIVICs form in the pericellular space between neighboring fat body cells.

In embryonic development and disease, epithelia can undergo an epithelial-to-mesenchymal transition (EMT). During this process, epithelial cells lose their apical-basal cell polarity as well as cell-cell and cell-BM adhesion to gain mesenchymal characteristics enabling them to migrate [15]. Some cancer cells can also undergo an epithelial-to-amoeboid transition (EAT) and use amoeboid cell migration to leave the tumor [16].

The *Drosophila* fat body undergoes fat body remodeling during metamorphosis at the early pupal stage at 4-14h after puparium formation (APF). Cells lose cell-cell and cell-BM adhesion and become individual cells that spread across the body within the hemolymph following head eversion [17, 18]. This process is induced by signaling through the steroid hormone Ecdysone and requires expression of the matrix metalloproteinases MMP1 and MMP2 [13, 17]. We recently discovered that at a later pupal stage, at 16h APF, fat body cells (FBCs) in the pupa are not passively floating in hemolypmph but are instead motile. They use swimming migration, an unusual subtype of amoeboid cell migration to respond to wounds [19, 20] and to patrol the pupa [21]. This suggests that FBCs must become migratory following fat body remodeling.

Overall, the larval fat body appears to have some similarities to epithelia. Both form cell layers through cell-cell and cell-BM adhesion. Yet cells in the fat body have a BM on each surface [22] while epithelia have a BM underlying only the basal surface [23]. Moreover, the cells in the fat body adhere to each other via CIVICs and are not known to have an apical-basal polarity. This raises the question of how CIVIC formation in the fat body is regulated and whether the apical-basal polarity machinery is involved.

Here we examine the epithelial apical-basal cell polarity network in the larval fat body and show that this tissue displays an apical-basal cell polarity. We find that aPKC is essential for cell-cell adhesion by regulating CIVIC formation in the larval fat body. We further show that apical-basal cell polarity and CIVICs are lost early during fat body remodeling which is regulated by Ecdysone signaling.

## Results

### The larval fat body tissue exhibits apical-basal cell polarity

To establish whether the fat body tissue in wandering third instar stage larvae has an apical-basal cell polarity, we performed antibody stainings for a range of classic polarity proteins known to localize to the apical (aPKC, Par-6, Crumbs) or basolateral domain (Dlg) in classic epithelia. Since it is not possible to image fat body cells from top to bottom at a high resolution due to their large size and light scattering issues, we mounted the fat body between two coverslips and imaged both sides of the tissue separately. To distinguish the two sides of the fat body (side (a) facing outwards towards the body wall and side (b) facing inwards towards the gut, Figure 1A on left), we took advantage of a morphological asymmetry noticeable in the larval fat body tissue architecture and only used the right sheet of the fat body for our experiments (see methods for more details). In addition to the immunostaining for particular polarity proteins, CAAX-GFP expression was used to visualize membranes to find the cell surface and lateral domains. We then quantified the mean intensities for CAAX-GFP and the antibody stain in several ROIs at the surface and at the lateral domain near the surface (shown in yellow and orange boxes, respectively in Figure 1A on right) on each side of the fat body for each animal (data in graphs paired for each tissue). These quantifications showed that while CAAX-GFP was always equally distributed on both surfaces and lateral domains (Suppl. Figure 1), aPKC, Par-6 and Crumbs consistently localized more strongly to the surface of side (a) with no difference in the lateral domains (Figure 1B-D’’). In contrast, Dlg localized more strongly to the surface on the opposite side, side (b), as well as to the lateral domain near side (b) (Figure 1E-E’’). A similar localization was also observed for another basolateral protein, Lgl, using a Lgl-GFP protein trap line (Figure 1F-F’’). Altogether, these results suggest that the larval fat body displays an apical-basal polarity with aPKC, Par-6 and Crumbs found on side (a), here forth referred to as apical side, and Dlg as well as Lgl on side (b), referred to as basal side, as well as spread along the basolateral domain.

**Figure 1.**
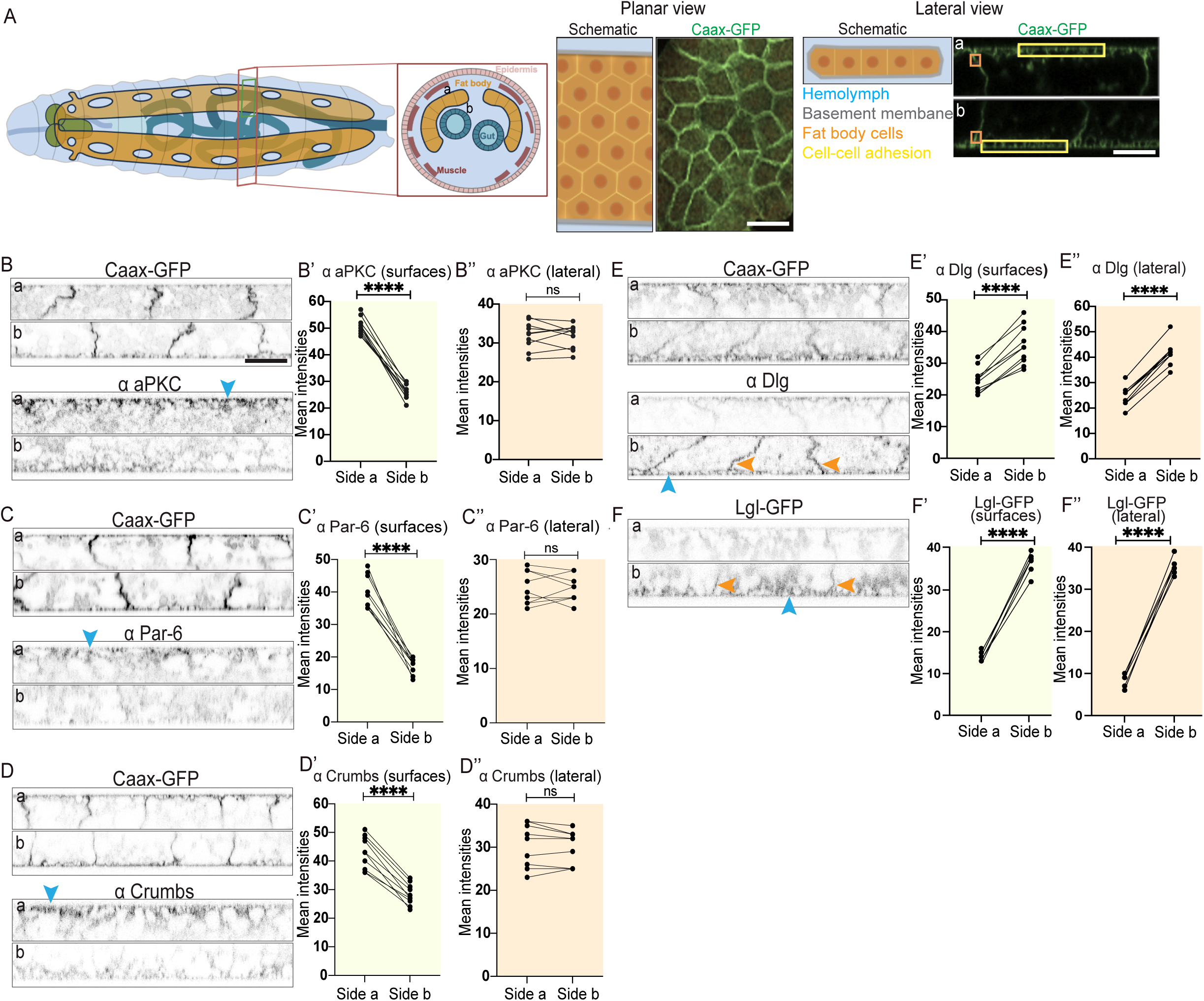
The larval fat body tissue exhibits apical-basal cell polarity. (A) Schematic of wandering third instar larva (dorsal view and cross section on left) showing location of fat body (orange, side a and b of fat body shown in cross section) in relation to brain (green), digestive system (blue), muscle (dark red) and epidermis (pink). Schematic and confocal images of CAAX-GFP-expressing fat body in planar view and lateral view (imaged from both sides; showing yellow surface ROIs and orange lateral ROIs used for intensity quantifications). (B-F’’) Confocal images of CAAX-GFP-expressing (B-E) or Lgl-GFP-expressing (F) larval fat body immunostained for aPKC (B), Par-6 (C), Crumbs (D) and Dlg (E; imaged from side a (top) and b (bottom), shown in lateral view, blue and orange arrowheads pointing at cell surfaces or lateral domains, respectively). Quantification of mean intensities of aPKC (B’, B’’), Par-6 (C’, C’’), Crumbs (D’, D’’), Dlg (E’, E’’) and Lgl-GFP (F’, F’’) on surface ROIs (‘, yellow background) or lateral ROIs (‘‘, orange background) on side a and b (mean of mean intensities from several ROIs at the surface or lateral domain of same tissue, data paired by tissue; n: 10 tissues, 3 surface or lateral ROIs per side (B’-E’ and B’’-E’’) and n: 6 tissues, 2 surface or lateral ROIs per side (F’, F’’)). Paired T-test, ****p<0.0001, ns p>0.05. Scale bars, 50µm (A - planar view image), 20µm (A - lateral view image and B-F)

### Cell-cell adhesion in the larval fat body does not require E-Cadherin but involves Collagen IV-Intercellular-Adhesion-Concentrations

Having discovered that the larval fat body tissue displays an apical-basal cell polarity, we wondered whether this polarity is involved in the regulation of cell-cell adhesion in the fat body, as in epithelia. Cell-cell adhesion in the fat body has been suggested to involve E-Cadherin-based adherens junctions [13]. However, whether E-cadherin is essential for cell-cell adhesion in the larval fat body, is not known. To investigate the role of E-Cadherin in cell-cell adhesion in the larval fat body further, we assessed the localization of E-Cadherin and Baz which are both known to localize to adherens junctions in the form of an apicolateral belt in many epithelia in *Drosophila* [3, 5, 6]. Our immunostainings using Baz and E-Cadherin antibodies overall resulted it a rather diffuse signal that labeled cell-cell vertices (marked with either CAAX-GFP or Lpp-Gal4+UAS-Myr-td-Tom), albeit relatively weakly and showed no clear belt-like apicolateral concentration (Figure 2A-B’). Our intensity quantifications revealed that both Baz and E-Cad localized more strongly to the apical surface and apicolateral domain than to the basal and basolateral domain, respectively (Figure 2A’-A’’’, B’-B’’’).

**Figure 2.**
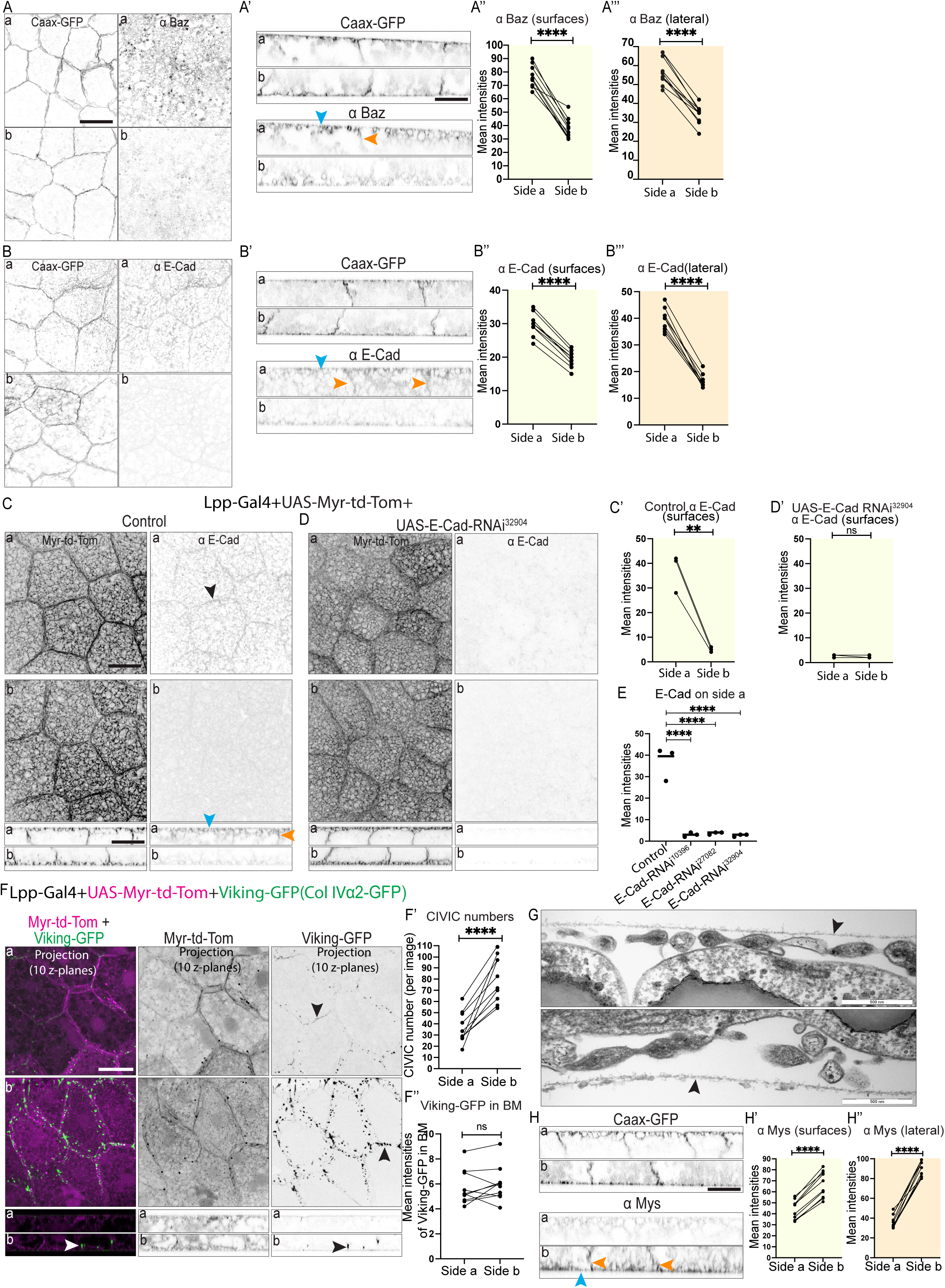
Cell-cell adhesion in the larval fat body does not require E-Cadherin but involves Collagen IV-Intercellular-Adhesion-Concentrations (A-B’’’) Confocal images of CAAX-GFP-expressing larval fat body immunostained for Baz (A, A’), and E-Cadherin (B, B’; side a (top) and b (bottom) in planar view (A, B) and lateral view (A’, B’), blue and orange arrowheads pointing at cell surfaces or lateral domains, respectively). Quantification of mean intensities of Baz (A’’, A’’’) and E-Cadherin (B’’, B’’’) on surfaces (‘‘, yellow) or lateral domains (‘’’, orange) on side a and b (mean of mean intensities from several ROIs, data paired by tissue; n: 10 tissues, 10 or 5 surface or lateral ROIs per side for Baz or E-Cadherin, respectively (A’’-B’’ and A’’’-B’’’)). Paired T-test, ****p<0.0001. (C-E) Confocal images of larval fat body expressing Lpp-Gal4+UAS-Myr-td-Tomato +control (C) or +UAS-E-Cadherin RNAi^32904^ (D) immunostained for E-Cadherin (side a and b shown in planar (top) and lateral views (bottom), black arrowhead pointing at cell-cell vertex, blue and orange arrowheads pointing at surface or lateral domain, respectively). Quantification of mean intensity of E-Cadherin for control (C’) or UAS-E-Cadherin RNAi^32904^ (D’) on surfaces on side a and b (mean of mean intensities from several ROIs, data paired by tissue; n: 3 tissues, 3 surface ROIs per side). Unpaired T-test, ****p<0.0001. Quantification of mean intensities of E-Cadherin for control, UAS-E-Cadherin RNAi^32904^ (from C’ and D’), UAS-E-Cadherin RNAi^103962^ and UAS-E-Cadherin RNAi^27082^ (from Suppl. Fig. 2B’, C’) shown for side a (E). Ordinary one-way multiple comparisons ANOVA, ****p<0.0001. (F-F’’) Confocal images of larval fat body expressing Lpp-Gal4+UAS-Myr-td-Tomato+Viking-GFP (F; side a and b shown in planar (top) and lateral views (bottom), black and white arrowheads pointing at CIVICs at cell-cell vertices; merge and single channels of Z projection of 10 Z planes at 2.5-5μm from cell surface). Quantification of CIVIC numbers per image (F’, using thresholded Z projection images of 10 Z planes of Viking-GFP channel (2.5-5μm from cell surface), n: 10 tissues, 10 Z projection images per tissue and side, data paired by tissue) and mean intensity of Viking-GFP in the basement membrane (F’’, using surface ROI in lateral view, data paired by tissue) on side a and b (n: 10 tissues, 3 lateral ROIs per tissue and side). Paired T-test, ****p<0.0001, ns p>0.05. (G) Transmission electron microscopy images of wild type larval fat body showing the basement membrane (arrowheads) near the cell surface on opposite sides of the tissue. (H-H’’) Confocal images of CAAX-GFP-expressing larval fat body immunostained for Mys (H, side a (top) and b (bottom), lateral view, blue and orange arrowheads pointing at cell surface or lateral domain, respectively). Quantification of mean intensity of Mys on surfaces (‘, yellow) or lateral domain (‘’, orange) on side a and b (mean of mean intensities from several ROIs, data paired by tissue; n: 10 tissues, 6 surface ROIs (H’) or 10 lateral ROIs (H’’) per side). Paired T-test, ****p<0.0001. Scale bars, 20µm (A-D, F, H), 500nm (G)

Next, we tested if E-Cadherin knockdown in the larval fat body is sufficient to cause cell-cell dissociation. Lpp-Gal4 was used to drive expression of UAS-E-Cad RNAi together with a membrane marker (UAS-Myr-td-Tom) specifically in fat body throughout larval stages. The third instar larval fat body was then immunostained for E-Cadherin to assess knockdown efficiency. E-Cad RNAi using three different independent RNAi constructs did not result in any cell-cell dissociation despite resulting in strong reduction in E-Cadherin staining demonstrating the efficiency of the RNAi knockdowns (Figure 2C-E, Suppl. Figure 2). This suggests that E-Cad knockdown is not sufficient to cause cell-cell dissociation in the larval fat body.

Apart from adherens junctions, Integrin-binding to pericellular CIVICs has been suggested to regulate cell-cell adhesion in the fat body [14]. Col IVα1 RNAi, Col IVα2 RNAi or Integrin β RNAi causes moderate cell-cell-dissociation of FBCs particularly on tricellular vertices [14], strongly suggesting that CIVICs mediate cell-cell adhesion in the fat body. Hence, we decided to study the distribution of CIVICs in the larval fat body along the lateral domain by looking at Viking-GFP expressed under its endogenous promoter (using a GFP protein trap in the Col IV α2 chain called Viking) as well as Lpp-Gal4+UAS-Myr-td-Tom to visualize membranes. We then quantified the number of CIVICs in the lateral domain near the apical and basal surfaces (using Z projections of 10 Z-layers at 2.5-5μm from cell surface on side a or b). As reported before [14], we saw CIVICs as punctae spread along the cell-cell vertices of FBCs (Figure 2F, note that the broader distribution of CIVICs along cell-cell vertices Z projection due to vertices sloping along Z axis). However, we found fewer CIVICs present at the lateral domain near the apical surface (Figure 2F-F’ side a). Our data, together with the findings from a previous study [14], suggests that cell-cell adhesion in the larval fat body is mainly mediated by CIVICs rather than by adherens junctions which are not essential for cell-cell adhesion.

Our discovery that the larval fat body displays apical-basal polarity raised the question of whether the basement membranes on the basal and apical surface [22] are the same. To assess this, we revisited our data from the CIVICs experiment since Viking-GFP labels CIVICs as well as basement membranes. When we compared mean intensities on both surfaces of the fat body, there was no significant difference between the two sides (Figure 2F’’). Moreover, electron microscopy on both surfaces of dissected fat body also did not reveal any obvious difference in basement membrane thickness (Figure 2G). Next, we assessed the localization of Mys, the main Integrin β subunit in flies, on the two surfaces of the fat body by immunostaining. Surprisingly, this revealed that, in contrast to Col IV, Integrin was more concentrated on the basal side as well as the basolateral domain of the fat body (Figure 2H-H’’) where CIVICs are found. Together, this suggests that the basement membrane is likely of similar thickness on both sides of the larval fat body.

However, the higher concentration of Integrin on the basal side suggests that there might be some differences between the basement membranes on the two surfaces.

### Apical-basal polarity regulates Collagen IV-dependent cell-cell adhesion in the larval fat body

Our discovery that the larval fat body tissue exhibits apical-basal polarity opened intriguing questions about the functional importance of this polarity. In epithelia apical-basal polarity proteins are known to regulate cell-cell adhesion via E-Cadherin-based adherens junctions [2]. To explore the role of apical-basal polarity in the fat body, we next investigated the effects of knocking down the polarity proteins aPKC, Crumbs, Scribble and Lgl in the fat body. To do this, we first imaged DAPI-stained fat body from wandering third instar larvae expressing UAS-aPKC-RNAi^34332^, UAS-Crumbs-RNAi^39177^, and UAS-Scribble-RNAi^105412^ together with a membrane marker (UAS-Myr-td-Tom) under the control of an early fat body driver Lpp-Gal4. Knocking down of *aPKC*, c*rumbs* and *scribbl*e resulted in partial cell-cell dissociation never seen in the control (Figure 3A-D, bicellular and tricellular gaps shown with white or yellow arrowheads, respectively). Moreover, knocking down *aPKC* and *scribble* using a second RNAi line (UAS-aPKC-RNAi^105624^, UAS-Scribble-RNAi^35748^), *crumbs* using two additional RNAi lines (UAS-Crumbs-RNAi^34999^ and UAS-Crumbs-RNAi^330135^) and *lgl* (UAS-Lgl-RNAi^109604^) also resulted in partial dissociation of cells (Suppl. Figure 3), validating our results further.

**Figure 3.**
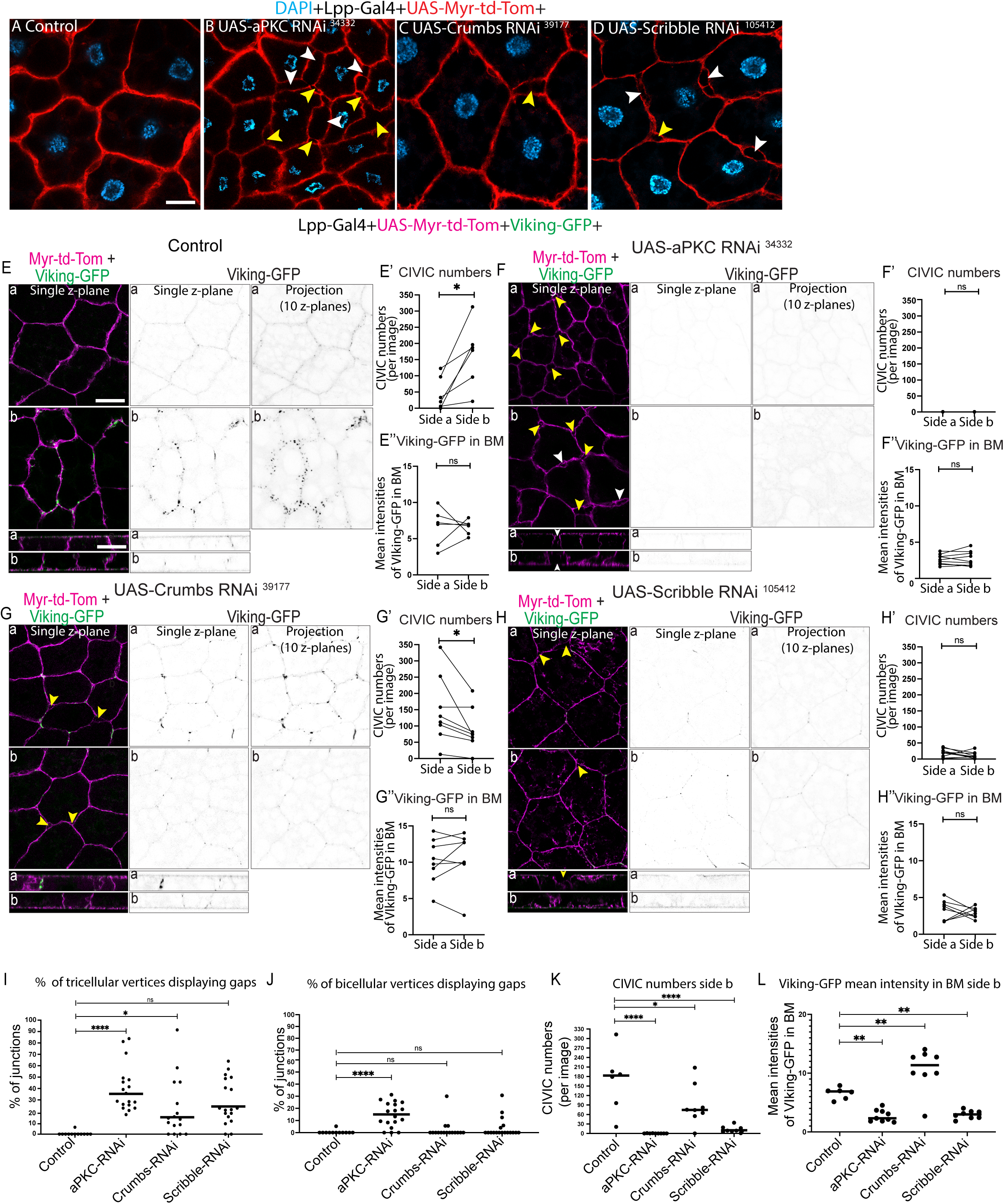
Apical-basal polarity regulates Collagen IV-dependent cell-cell adhesion in the larval fat body (A-D) Confocal single Z plane images of DAPI-stained, larval fat body expressing Lpp-Gal4+UAS-Myr-td-Tomato +control (A), UAS-aPKC RNAi^34332^ (B), UAS-Crumbs RNAi^39117^ (C) and UAS-Scribble RNAi^105412^ (D; yellow or white arrowheads showing gaps at tricellular or bicellular cell-cell vertices, respectively). (E-L) Confocal images of larval fat body expressing Lpp-Gal4+UAS-Myr-td-Tomato+Viking-GFP +control (E), +UAS-aPKC RNAi^34332^ (F), +UAS-Crumbs RNAi^39117^ (G) and +UAS-Scribble RNAi^105412^ (H; showing images of merged channels (single Z plane) and Viking-GFP channel (single Z plane and Z projection of 10 layers 2.5-5μm from cell surface); yellow or white arrow showing gaps at tricellular or bicellular vertices, respectively). Quantification of percentage of tricellular or bicellular cell-cell vertices containing gaps per image (I, J, respectively) from E-H (n: 6 images (control), 9 images (UAS-aPKC RNAi^34332^), 8 images (UAS-Crumbs RNAi^39117^), and 8 images (UAS-Scribble RNAi^10541^), each from different pupae). Ordinary one-way multiple comparisons ANOVA, ****p<0.0001. Quantification of mean CIVIC numbers (E’, F’, G’, H’; from thresholded Z projection images of 10 Z planes of Viking-GFP channel (2.5-5μm from cell surface), n: 10 tissues, 10 Z projection images per tissue and side, data paired by tissue) and mean intensity of Viking-GFP in the basement membrane (E’’, F’’, G’’, H’’; using surface ROIs in lateral view, data paired by tissue; n: 6 pupae (control), 9 pupae (UAS-aPKC RNAi^34332^), 8 pupae (UAS-Crumbs RNAi^39117^), and 8 pupae (UAS-Scribble RNAi^105412^), 1 ROIs per side for each tissue) on side a and b. Paired T-test, ****p<0.0001. Quantification of CIVIC numbers (K) and mean intensity of Viking-GFP in the basement membrane (L) for control, UAS-aPKC RNAi^34332^, UAS-Crumbs RNAi^39117^ and UAS-Scribble RNAi^105412^ on side b. Ordinary one-way multiple comparisons ANOVA, ****p<0.0001. Scale bars, 20µm (A-D, E-H)

To quantify the extent of cell-cell dissociation and to assess if these defects are due to aberrant CIVICs-mediated cell-cell adhesion, we next expressed UAS-aPKC-RNAi^34332^, UAS-Crumbs-RNAi^39177^, and UAS-Scribble-RNAi^105412^ with Lpp-Gal4 alongside with UAS-Myr-td-Tom for membrane labeling and Viking-GFP to quantify CIVIC numbers. We found that aPKC-RNAi^34332^, Crumbs-RNAi^39177^, and Scribble-RNAi^105412^ resulted in 35%, 15% and 25% of tricellular vertices showing gaps, respectively (Figure 3E, F, G, H and I). aPKC-RNAi^34332^ also resulted in 15% of bicellular vertices having gaps (figure 3E, F, G, H and J).

Quantifications of CIVIC numbers on both cell surfaces revealed that the asymmetric localization of CIVICs, with higher numbers seen basolaterally than apicolaterally in the control (Figure 3E, E’), as seen before, was disrupted upon aPKC-RNAi^34332^, Crumbs-RNAi^39177^, and Scribble-RNAi^105412^ (Figure 3F, F’, G, G’, H and H’). Strikingly, aPKC and Scribble knockdown resulted in near complete or strong loss of CIVICs on both sides, respectively (Figure 3F, F’, H, H’ and K) as well as a reduction in Viking-GFP signal in the basement membranes (Figure 3F’’, H’’ and L). In contrast, Crumbs knockdown resulted in a redistribution of CIVICs along the lateral domain, with higher numbers of CIVICs on the apicolateral than on the basolateral region (Figure 3G and G’, K). Moreover, it increased the Viking-GFP signal in the basement membranes (Figure 3G’’, L).

Together, our results suggest that polarity proteins aPKC, Lgl, Scrib and Crumbs play an important role in regulating cell-cell adhesion and tissue organization in the larval fat body tissue. aPKC, in particular, plays a key role in regulating cell-cell adhesion by mediating CIVIC formation.

### Fat body cells dissociate during Ecdysone-regulated fat body remodeling to initiate amoeboid swimming migration

Having established that the larval fat body displays an apical-basal cell polarity that regulates an unusual, CIVIC-mediated cell-cell adhesion mechanism, we next wanted to investigate how cell-cell dissociation during fat body remodeling (FBR) is regulated. FBR happens around 4-14h APF [17, 18] (Figure 4A). Having recently discovered that FBCs are migratory in 16h APF pupae [19, 20], we suspected that FBCs become migratory following FBR. Indeed, we saw this when we imaged fat body remodeling *in vivo*. FBCs (nuclei marked) initially remained close to each other within the two lateral sheets and then moved slightly apart from each other (Figure 4B1 and B2, respectively, Movie 1). Soon after the rear retraction of the animal and head eversion, cells spread across the body (Figure 4B3, Movie 1) and started migrating as shown by the gradual increase in cell speed before plateauing 7h after head eversion (Figure 4B4, B5 and C, Movie 1).

**Figure 4.**
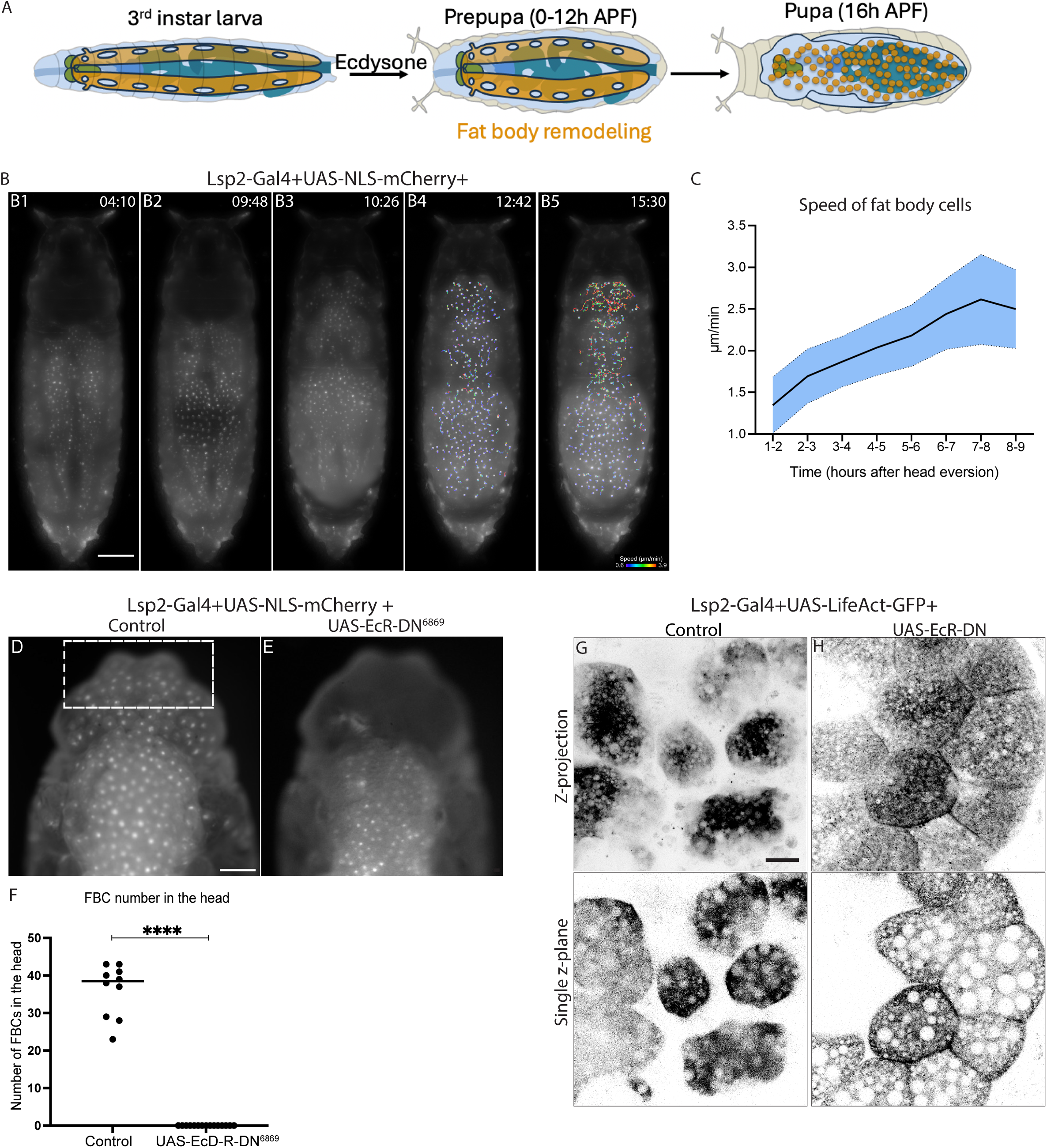
Fat body cells dissociate during Ecdysone-regulated fat body remodeling to initiate amoeboid swimming cell migration (A) Schematic showing fat body morphology before, during and after fat body remodeling (third instar larva, prepupa and 16h APF pupa). (B-C) Widefield time-lapse images of the dorsal view of a pupa expressing Lsp2-Gal4+UAS-NLS-mCherry (B1-B5, pupal age 4h APF at start of movie, imaged at room temperature). Time in hours: min. Migration tracks starting 1h after head eversion (tracks of minimum length of 90min): color-coded according to current speed (B4-B5). Quantification of mean current speed of FBCs in head and thorax over time (C; n: 357 tracks from 6 pupae). See Movie 1. (D-F) Widefield images of the dorsal view of head and thorax region of 16h APF pupae expressing Lsp2-Gal4+UAS-NLS-mCherry +control (D) or +UAS-EcRDN^6869^ (E). Quantification of number of FBCs in front of head (F; n: 10pupae (control), 15 pupae (UAS-EcRDN^6869^); cells counted in front half of head (dotted rectangle in D)). Mann-Whitney test, ****p<0.0001. (G-H) Confocal images of fat body expressing Lsp2-Gal4+UAS-LifeAct-GFP +control (G), UAS-EcR-DN^6869^ (H, Z projection top and single Z plane bottom). Scale bars, 300µm (B), 150µm (D-E), 20µm (G-H)

FBR was strongly blocked when we expressed a dominant-negative version of Ecdysone receptor (UAS-EcR-DN) together with the nuclear marker UAS-NLS-mCherry using Lsp-Gal4. While FBCs had moved into the head of control pupae after completion of FBR at 16h APF, the fat body in EcR-DN-expressing pupae remained as sheets in the thorax and abdomen, and no individual cells could be seen in the head (Figure 4D-F). Moreover, *in vivo* live imaging of Lsp-Gal4+UAS-Myr-td-Tomato+UAS-EcR-DN-expressing pupae at 16h APF further showed that the cells remained closely attached in the dorsal abdomen and thorax while control cells were seen as individual migratory cells (Figure 4G-H). This shows that Ecdysone signaling is essential for cell-cell-dissociation during FBR, as shown before [17, 24].

### Ecdysone regulates cell-cell dissociation through the loss of apical-basal cell polarity and CIVICs during fat body remodeling

Having found that apical-basal cell polarity regulates cell-cell adhesion in the larval fat body, we next wanted to see whether apical-basal polarity is lost during FBR before cells dissociate, as during classic EMT. For this, we imaged fat body tissues from 3h APF-old pupae expressing the membrane marker Ubi-CAAX-GFP immunostained for aPKC, Par-6, Crumbs, Baz or Dlg. This showed that the asymmetric localization of aPKC, Par-6, Crumbs and Baz to the apical surface or Dlg to the basal surface that we saw in the larval fat body (Figure 1B’-E’ and Figure 2A’’), was lost at 3hr APF for all these polarity proteins (Figure 5A-E’). This suggests that apical-basal cell polarity is lost early during fat body remodeling when the cells are still attached to each other.

**Figure 5.**
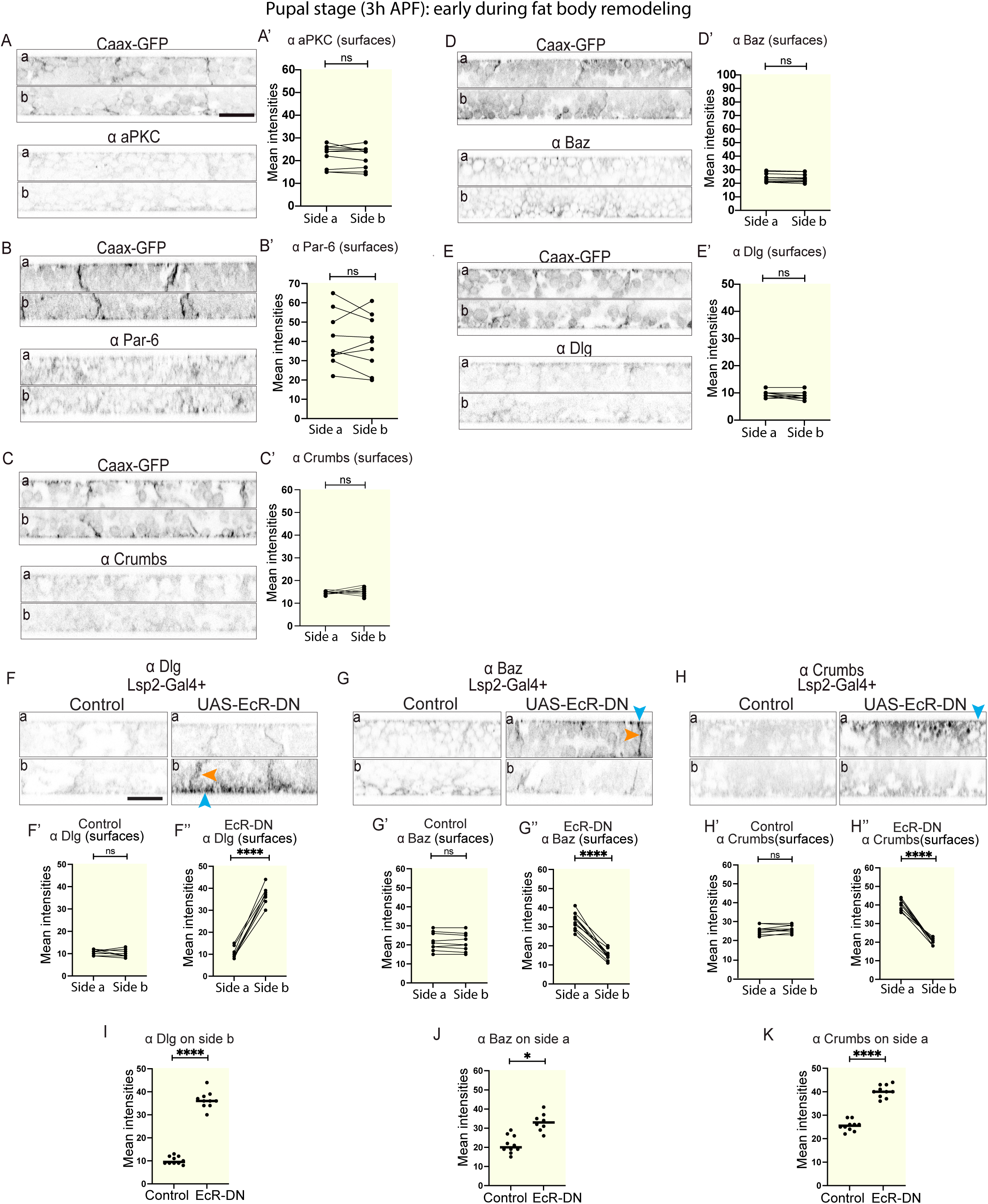
Ecdysone regulates the loss of apical-basal cell polarity during fat body remodeling (A-E’) Confocal images of fat body from CAAX-GFP-expressing, 3hAPF pupae immunostained for aPKC (A), Par-6 (B), Crumbs (C), Baz (D) and Dlg (E), imaged on sides a (top) and b (bottom), shown in lateral view. Quantification of mean intensities of aPKC (A’), Par-6 (B’), Crumbs (C’), Baz (D’) and Dlg (E’; at the surfaces, paired for each fat body tissue, n: 10 tissues, 3 surface ROIs per tissue per side). Paired T-test, ns p>0.05. (F-K) Confocal images of fat body from 3hAPF pupae expressing Lsp2-Gal4 +control or +UAS-EcRDN^6869^ (F, G, H, left and right, respectively) immunostained for Dlg (F), Baz (G) and Crumbs (H, side a (top) and b (bottom), lateral view, blue and orange arrowheads pointing at cell surface or lateral domain, respectively). Quantification of mean intensities of Dlg (F’-F’’), Baz (G’-G’’) and Crumbs (H’-H’’) on surfaces on side a and b for control or UAS-EcRDN^6869^ (mean of mean intensities from several ROIs, data paired by tissue; n: 10 tissues, 3 surface ROIs per tissue per side). Paired T-test, ****p<0.0001. Quantification of mean intensities of Dlg (I), Baz (J) and Crumbs (K), at the surface for control and UAS-EcRDN^6869^ on side b (I) or side a (J, K). Unpaired T-test, ****p<0.0001. Scale bars, 20µm (A-E, F-H)

Next, we wanted to investigate whether Ecdysone signaling regulates this loss of polarity. Immunostaining for Dlg, Baz and Crumbs showed that in pupae expressing Lsp2-Gal4+UAS-NLS-mCherry+UAS-EcR-DN, Dlg remained concentrated on the basal side, and Baz and Crumbs remained concentrated on the apical side (Figure 5F-K), similar to what we saw in larval fat body (Figure 1D’, E’ and Figure 2A’’), while their asymmetric localization was lost in the control pupae (Figure 5F-F’, G-G’, H-H’). This suggests that Ecdysone signaling in the fat body regulates the loss of apical-basal cell polarity in the fat body early during FBR.

Since cell-cell-adhesion in the third instar larval fat body is mediated by CIVICs [14], we assessed next whether CIVICs get lost from cell-cell vertices during FBR in wild type. Imaging of fat body from 3h APF pupae expressing Lpp-Gal4+UAS-Myr-td-Tomato+Viking-GFP showed that CIVIC numbers were very low in the WT fat body (Figure 6A, A’), much lower than the numbers seen in the WT fat body of third instar larvae (Figure 2F, F’). This shows that CIVICs are lost early during FBR. In contrast, animals expressing UAS-EcR-DN, failed to lose CIVICs and had much larger numbers of CIVICs than the control (Figure 6B, B’ and C), suggesting that Ecdysone signaling is needed for the loss of CIVICs from the cell-cell vertices early during FBR.

**Figure 6.**
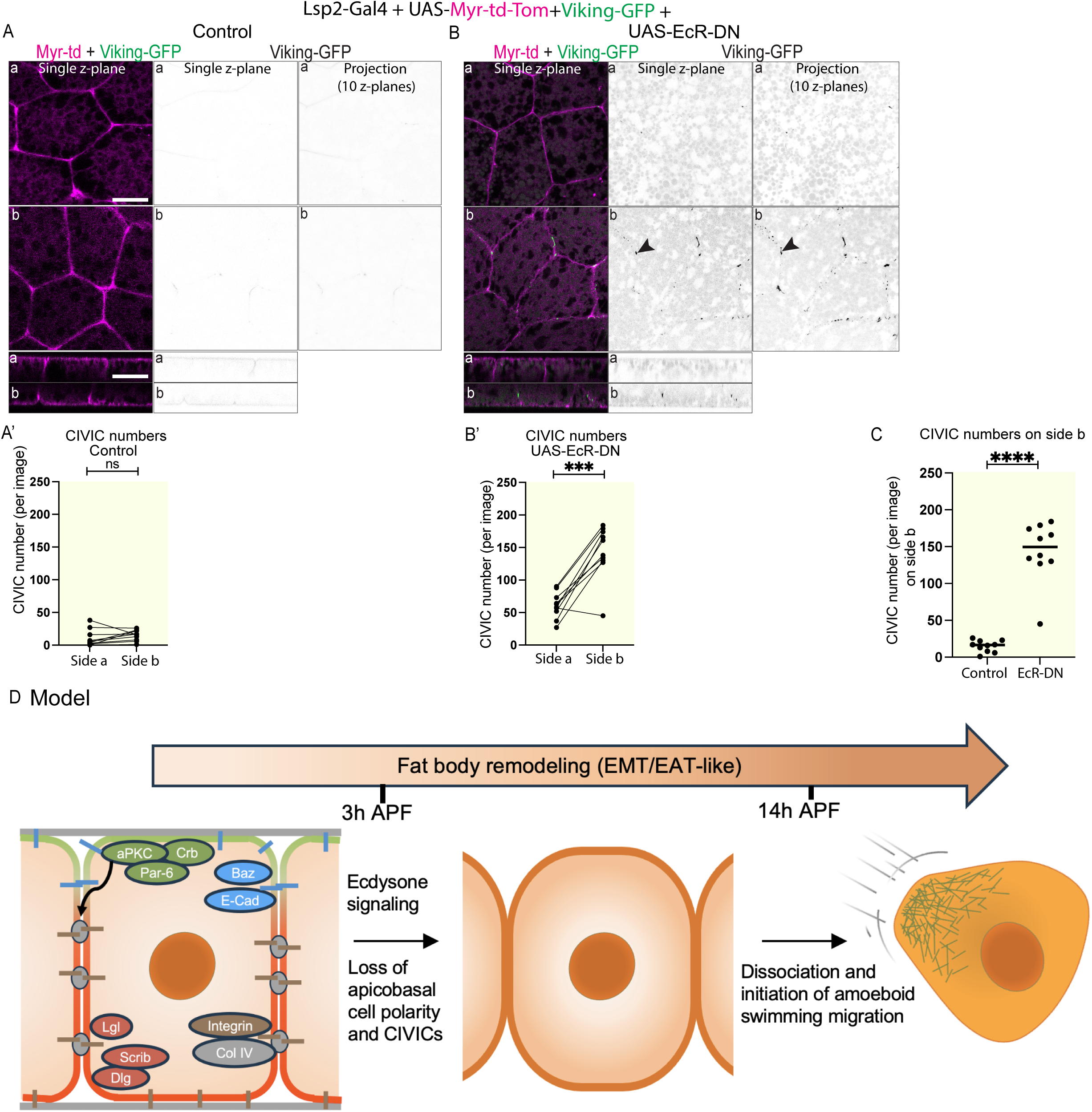
Ecdysone regulates cell-cell dissociation during fat body remodeling through the loss of CIVICs (A-C) Confocal images of fat body from 3h APF pupae expressing Lpp-Gal4+UAS-Myr-td-Tomato+Viking-GFP +control (A) or +UAS-EcRDN^6869^ (B; side a and side b shown in planar and lateral view; merge of single Z plane (left) and Viking-GFP channel of single Z plane or of projection of 10 Z planes 2.5-5μm from cell surface (middle and right panels, respectively)). Quantification of mean Viking-GFP-positive CIVIC numbers (A’, B’, from thresholded Z projection images of 10 Z planes of Viking-GFP channel (2.5-5μm from cell surface), n: 10 tissues, 3 Z projection images per tissue and side, data paired by tissue). Paired T-test, ****p<0.0001. Quantification of CIVIC numbers for control and UAS-EcRDN^6869^ on side b (C). Unpaired T-test, ****p<0.0001. (D) Proposed model of fat body remodeling. The fat body in the third instar larva displays an apical-basal cell polarity which regulates Collagen IV-mediated cell-cell adhesion. Ecdysone induces fat body remodeling in the prepupa resulting in the loss of apical-basal polarity and CIVICs by 3h APF. Cells then dissociate and initiate amoeboid swimming migration in pupae around 14h APF. Scale bars, 20µm (A-B)

## Discussion

Apical-basal cell polarity is a key hallmark of epithelia, where it dictates their structure and function. One of its key roles is in regulating cell-cell adhesion. Despite being of non-epithelial nature, the mesoderm-derived adipose tissue in flies and humans is composed of tightly associated cells. The mechanisms that regulate adipose tissue architecture as well as the functional significance of this architecture are still mostly unknown. Here we show that the adipose tissue in flies, the fat body, previously not thought to be polarized, in fact displays an apical-basal cell polarity which is essential for cell-cell adhesion and tissue integrity (Figure 6D). Strikingly, in contrast to epithelia, the apical-basal polarity machinery in the fat body regulates cell-cell adhesion via Collagen IV rather than E-Cadherin. An interesting question that arises from this discovery is what the underlying mechanism is? Apical-basal polarity proteins usually determine the apical, apicolateral, basolateral and basal cell domains. These, in turn, might regulate the architecture of the cytoskeleton which could direct localized secretion of Col IV and/or transport of Integrin to specific sites in the lateral domain to initiate CIVIC formation. Interestingly, Col IV fibrils, which are of similar composition and structure to CIVICs, have been described to form at cell-cell interphases in *Drosophila* follicle cells, a type of somatic epithelial cells [25]. These Col IV fibrils form through directed Rab10-mediated secretion of Col IV into the pericellular space at the basal region of cell-cell interfaces, from where they are subsequently incorporated into the underlying BM [25]. A similar mechanism of directed Col IV transport into the lateral plasma membrane might also mediate CIVIC formation in the fat body.

Col IV-dependent cell-cell adhesion has so far only been reported for the fat body, but it might not be unique to this tissue. As mentioned above, Col IV fibrils that form in the pericellular space in follicle cells have a similar localization and ECM composition to CIVICs [25] and might hence be related structures. While these Col IV fibrils have not been reported to mediate cell-cell adhesion, it is tempting to speculate that they could contribute to cell-cell adhesion in the basolateral cell region of follicle cells before being deposited into the BM. Moreover, there are also some known examples where Integrin-binding to ECM can mediate cell-cell adhesion. For example, Integrins at myotendinous junctions are known to connect muscle cells to tendon cells through an intervening ECM [26].

It may be that the use of CIVICs for cell-cell adhesion is unique to tissues that secrete Collagen IV like the fat body. Interestingly, there are two populations of Col IV that FBCs [14] and follicle cells [25] produce, one secreted to the surface to form the BM and one secreted laterally to form Col IV concentrations in the pericellular space. Hence, it might be that CIVIC-dependent cell-cell adhesion and focal adhesion-mediated cell-basement membrane adhesion are part of an interlinked mechanism that has coexisted during evolution. Early in evolution apical-basal polarity might have regulated Integrin-binding to locally secreted Col IV-containing ECM which in turn could have mediated cell-cell adhesion at the same time as mediating cell-basement membrane adhesion. This could hence constitute an ancient mechanism enabling the evolution of multicellular animals from their unicellular ancestors. Both the integrin adhesion machinery [27] and the E-Cadherin adhesion machinery [28] have been reported to have an ancient origin predating the emergence of metazoan and might hence have evolved in parallel. In most tissues in extant animals, E-Cadherin-mediated adhesion might then have then taken a dominant role over Col IV-dependent cell-cell adhesion.

The apical-basal cell polarity that we observe in the larval fat body seems similar to the one in most classic epithelia. However, one key difference is that in the fat body there is a BM containing Col IV on both the apical and basal surfaces. In contrast, epithelia only have a Col IV-containing BM on the basal side, while they also often have an apical ECM devoid of Col IV. Interestingly though, despite the presence of a BM on both surfaces, we see Integrin concentrated only on the basal surface of the fat body. It may hence be, that the apical and basal BM, despite having a similar appearance by EM and containing similar levels of Col IV, might differ otherwise in their ECM composition with Laminin being a potential candidate since it can bind Integrin [29] and affect apical-basal polarity [30]. Further research is needed to reach a better molecular understanding of how similar apical-basal cell polarity and the apical and basal ECM/BM are in the fat body compared to epithelia.

What is the function of apical-basal cell polarity in the larval fat body? Our results show that the apical-basal polarity proteins aPKC, Crumbs, Lgl and Scribble are required for intercellular adhesion of fat body cells. aPKC, in particular, plays a key role here in mediating cell-cell adhesion by regulating CIVIC formation. Apart from this, it seems likely that apical-basal cell polarity also plays other roles in regulating cell function. It is tempting to speculate that some of the well-established functions of the fat body, including lipid uptake or release [31], antimicrobial peptide secretion to fight infection [32] or tumors [33] or secretion of other factors such as growth factors [34–36] might be mainly mediated via either the apical or the basal cell surface.

Apart from providing new insight into how adipose tissue architecture is regulated, our study also sheds new light into the process of fat body remodeling. We show Ecdysone signaling in the fat body induces cell-cell-dissociation by regulating the loss of apical-basal cell polarity and CIVICs (Figure 6D). Interestingly our new findings, together with our previous findings [21] show that fat body remodeling is followed by initiation of amoeboid swimming migration. FBR and EMT have several key features in common. First, in both cases there is an apical-basal cell polarity mediating cell-cell adhesion, albeit through different mechanisms, which is then lost during the process to induce cell-cell dissociation. Second, matrix metalloproteinases induce loss of cell-basement membrane adhesion [13, 17, 37]. Third, at the end of the process, in both cases cells become motile even though using different migration modes. FBCs use amoeboid swimming cell migration [21] whilst most cells undergoing EMT that have been studied so far use mesenchymal cell migration. However, some cancer cells undergo an epithelial-to-amoeboid-transition resulting in amoeboid cell migration [16]. Interestingly, during wound healing in mice, adipocytes have been shown to become migratory to invade into the wound bed, suggesting that they might also undergo an EMT-like process [38, 39]. It remains to be seen, if the adipose tissue in mammals might also display an apical-basal polarity.

Taking all of this into account, we propose that the remodeling of the *Drosophila* adipose tissue constitutes a novel category on the spectrum of epithelial-to-mesenchymal/amoeboid transition. This powerful genetic *in vivo* model system could be a valuable addition to the small set of commonly used EMT models which could be helpful to unravel the diverse mechanisms underlying EMT and EAT in health and disease.

## Materials and Methods

### Fly stocks and maintenance

*Drosophila* melanogaster stocks and crosses were maintained and performed on cornmeal molasses food at 25°C. Stocks obtained from Bloomington *Drosophila* Stock Center (NIH P40OD018537) were used in this study. The following lines were used in this paper: w^67^ as a control, Lsp2-Gal4 (BDSC: 6357), Lpp-Gal4 (gift from Pierre Leopold), Ubi-CAAX-GFP (DGRC: 109824), UAS-EcR-B1-DN (BDSC:6869), UAS-LifeAct-GFP [40], UAS-Myr-td-Tom (BDSC: 32221), UAS-NLS-mCherry (BDSC: 38424), Lgl-GFP (BDSC: 63183), UAS-aPKC-RNAi 1 (BDSC: 34332) and 2 (BDSC:105624), UAS-Scribble-RNAi 1 (BDSC: 35748) and 2 (BDSC: 105412), UAS-Crumbs-RNAi 1 (BDSC: 34999), 2 (VDRC: GD-39177), 3 (VDRC: shRNA-330135) and 4 (BDSC: 40869), UAS-Lgl-RNAi (VDRC: KK-109604), UAS-ECad-RNAi (VDRC: KK-103962), 2 (VDRC: GD-27082) and 3 (BDSC: 32904) and Viking-GFP (DGRC: 110626). We used FlyBase to find information on phenotypes/function/stocks/gene expression (etc.).

### Fat body dissections

Wandering third instar larvae were dissected on a sylgard-coated depression dish. Animals were placed on their dorsal side and pinned by the tail and mouth hooks. Using spring-scissors a horizontal incision was made in the posterior end of the larva, followed by a vertical cut along the dorsal midline towards the rostral end of the larva. Then a horizontal cut was made left and right of the pin at the rostrum of the animal. The flaps were then pinned in a clockwise order to ensure that the animal’s body is stretched both horizontally and vertically. The animal was fixed with 4% Paraformaldehyde for 30 minutes, to allow organs to float and facilitate organ removal, including fat body tissues. For fat body dissections, the trachea along with the gut were first removed, ensuring the fat bodies were kept along the sides of the animal. An incision on the anterior end of the right side of the fat body was then made. The right halves of fat body tissues were then placed in 96-well plates and washed twice with PBS. The dissected tissues were sometimes stored in PBS for up to 5 days at 4°C. This dissection method was also used to dissect fat body tissues from 3h APF pupae.

### Immunohistochemistry

Dissected fat body tissues were fixed in PBS containing 4% Paraformaldehyde for 30 minutes, permeabilized in PBS containing 1% Trition-X-100 at room temperature and blocked in PBT with 4% fetal bovine serum. The dissected tissues were incubated overnight at 4°C with primary antibodies mouse anti-Crb (1:50; DSHB AB_528181 (cq4)), mouse anti-Dlg1 (1:20; BSHB AB_528203 (4f3)), rabbit anti-Baz (1:2000; a gift from Andreas Wodarz, University of Cologne), guinea pig anti-Par-6 (1:500; [41]), rabbit anti-aPKC (1:500; SAB4502380, Sigma), rat anti-DE-Cadherin (1:20; DSHB AB_528120 (E-CAD2) and mouse anti-Mys (1:20; DSHB AB_528310 (cf.6g11)) diluted in PBT. After three washes in PBS, the dissected tissues were incubated with secondary antibody, anti-rat 567nm (1:200; abCAM), anti-mouse 567nm (1:200; abCAM), anti-guinea pig 567nm (1:200; abCAM), anti-rabbit 567nm (1:200; abCAM), for 2 hours at room temperature. Fixed and stained samples were mounted on DAPI-vectashield and prepared for imaging.

### Microscopy

#### Imaging setup to image dissected fat body

Stained right halves of fat body tissues dissected from different animals were mounted between two coverslips. Side (a) and side (b) of the right half of the fat body could be identified the following way: The fat body sheet has several round gaps in the tissue. The tissue is wider (∼4-7 FBCs wide) on one side of the gaps than on the other side (∼1-2 FBCs wide, see Figure 1A). For imaging side (a), the fat body sheet was oriented with the anterior end upwards and the posterior end downwards ensuring that the thicker side was pointing left and the thinner side right. To image side (b) the cover glass was flipped over so that the anterior end pointed upwards and the posterior end downwards, and the thinner side was located to the left, and the thicker side to the right.

#### Imaging setup to image dissected pupae

For imaging done on pupae, animals were kept at 25°C. Pupae were marked at the white pre-pupa stage (0h AFP) and dissected at 16h APF by removing the pupal case [42] and placed on a coverslip on their dorsal side for imaging.

#### Microscopy

Microscope images were collected on a Zeiss LSM 980 microscope using 63x objectives at 0.25μm step size except for Figure 4B, D-E and Movie 1 which were collected on a Zeiss Cell discovery 7 widefield microscope with a 5X lens at 0.5 magnification. Movie 1 was acquired for 19h24min with a time interval of 2min at room temperature. Nuclear tracking in Movie 1 was done as described before [21]. Average migration speed per pupa was obtained by averaging the mean speed of all individual tracks at various 1h time intervals (e.g. 1-2h, 2-3h etc. post head eversion). Tracks (of a minimum tracking length of 90min) were color-coded based on their current speed over time in IMARIS. Tracks are shown in images as whole tracks dragon tail tracks.

Images and movies of confocal movies were generated with Fiji ImageJ to create Z-projections, Z-sections and orthogonal view images. We used the same brightness and contrast adjustment for control and experimental conditions. Movies and images were organized and annotated with VSDC Video Editor and Abode Illustrator.

### Electron microscopy

Dissected fat body tissues from wandering third instar stage larvae were fixed in 2% Paraformaldehyde, 1.5% (v/v) glutaraldehyde in 0.1 M cacodylate. Following 0.1M cacodylate washes, fat body tissues were fixed in 1% osmium tetroxide and 1.5% K3[Fe(CN6)] for 1 hour at room temperature and rinsed with ddH_2_O. After osmium-ferricyanide staining, the tissues were treated with 1% thiocarbohydrazide (TCH) for 20 minutes at room temperature, stained with 2% osmium tetroxide for 30 minutes at 4°C, washed three times with ddH_2_O and stained with 1% aqueous uranyl acetate overnight at 4°C. Next, the tissues were incubated with freshly made lead aspartate solution for 30 minutes at 60°C. After being rinsed with buffer and gradually dehydrated with increasing concentrations of ethanol (70%, 90%, 100%), the tissues were infiltrated with a graded series of EPON. The tissues were then placed in blocks, polymerized in 100% resin and baked overnight in 60°C. Electron microscopy was performed with a Tecnai G2 Spirit transmission electron microscope (FEI) equipped with a Morada charge-coupled device camera (Olympus Soft Imaging Systems).

### Image analysis

#### CIVIC numbers

CIVICs quantifications were performed on the maximum projection of 10 layers within the Z-stack (2.5-5μm from cell surface). A threshold rage of 43-255 was applied in Fiji and particles with sizes from 0.2-infinity (pixel^˄^2) were identified and analyzed.

#### Analysis of bicellular or tricellular gaps

The number of bicellular or tricellular cell-cell vertices with or without gaps was counted manually using the cell counter plugin on FIJI-ImageJ at 7.5-20μm from cell surface to calculate the percentage of vertices containing gaps.

#### Mean intensity analysis at cell surfaces and lateral sides

Mean fluorescence intensities were measured using ROIs (one for surface measurements (46.69×3.29μm (wide/high)), shown in yellow in Figure 1A and one for lateral measurements (3.62×4.60μm (wide/high)), shown in orange in Figure 1A). The mean of the mean intensities of several regions imaged on the same side (either side a or b) of the same fat body tissue was calculated for each fat body tissue and shown paired per tissue in the graphs.

#### Fat body cell counts in pupal head

Fat body cells in the head of pupae (Figure 4D-E) were counted manually in the front half of the head of the pupae using the cell counter plugin on FIJI-ImageJ.

### Statistics

Statistics were performed using GraphPad Prism 9. Statistical tests used in each experiment are indicated in the relevant figure legends. P<0.05 was set as the significance threshold. In scatter dot plots, the line in the middle indicates the median.

## Acknowledgements

We would like to thank the Pichaud, Amoyel, Fernandes and Mao labs for critical discussions and the VDRC and BDSC for *Drosophila* stocks. Thanks to Franck Pichaud for reading the manuscript and to the UCL Biosciences Imaging Facility for imaging help. We thank Virginia Silio for help with image analysis. This work was funded by the Wellcome Trust and Royal Society Sir Henry Dale fellowship granted to A.F. (215431/Z/19/Z).

## Author contributions

Experimental design, data collection and analysis for all the experiments in Figures 1-6 and all the supplementary Figures was done by J.A. with the exception of Figure 5B and C and Movie 1 which were done by C.A. B.H.L. generated the animals used for Figure 3G-L. A.F. designed the study and J.A. and A.F. wrote the manuscript.

