## Supplemental Information for "Apical-basal polarity regulates Collagen IV-dependent cell-cell adhesion in the *Drosophila* adipose tissue"

### Supplementary legends

#### Supplementary movie legends

**Movie 1 – Fat body cells dissociate during fat body remodeling to initiate cell migration**

Widefield time-lapse series of the dorsal view of a pupa expressing Lsp2-Gal4+UAS-NLS-mCherry (pupal age 4h APF at start of movie, imaged at room temperature). Time in hours: min. Migration tracks starting 1h after head eversion (only showing tracks of minimum length of 90min): color-coded according to current speed. See Figure 4B.

#### Supplementary figure legends

**Supplementary Figure 1 – The larval fat body tissue exhibits apical-basal cell polarity – related to Figure 1B-E**

(A-D'') Confocal images of CAAX-GFP-expressing larval fat body immunostained for aPKC (A), Par-6 (B), Crumbs (C) and Dlg (D; imaged from side a (top) and b (bottom), merged channels shown in lateral view, CAAX-GFP in green and antibody stain in magenta).

Quantification of mean intensities of CAAX-GFP on surface ROIs (', yellow background) or lateral ROIs ('', orange background) on side a and b (mean of mean intensities from several ROIs at the surface or lateral domain of same tissue, data paired by tissue, quantifications using same samples as in Figure 1B-E; n: 10 tissues, 3 surface or lateral ROIs per side (B'-E' and B''-E'') and n: 6 tissues, 2 surface or lateral ROIs per side (F', F'')). Paired T-test, ns $p>0.05$.

Scale bars, 20µm (A-D)

**Supplementary Figure 2 – Cell-cell adhesion in larval fat body does not require E-Cadherin – related to Figure 2C-E**

(A-C'') Confocal images of larval fat body expressing Lpp-Gal4+UAS-Myr-td-Tomato +control (A), +UAS-E-Cadherin RNAi$^{103962}$ (B) or +UAS-E-Cadherin RNAi$^{27082}$ (C) immunostained for E-Cadherin (side a and b shown in planar (top) and lateral views (bottom)).

Quantification of mean intensity of E-Cadherin for control (A’), UAS-E-Cadherin RNAi[103962] (B’) or UAS-E-Cadherin RNAi[27082] (C’) on surfaces on side a and b (mean of mean intensities from several ROIs, data paired by tissue; n: 3 tissues, 3 surface ROIs per side). Unpaired T-test, ****$p<0.0001$, ns $p>0.05$.

Scale bars, 20µm (A-C)

**Supplementary Figure 3 – Apical-basal cell polarity is needed for Collagen-IV-dependent cell-cell adhesion – related to Figure 2A-D**

(A-F) Confocal single Z plane images of DAPI-stained, larval fat body expressing Lpp-Gal4+UAS-Myr-td-Tomato +control (A), UAS-aPKC RNAi[105624] (B), UAS-Crumbs RNAi[34999] (C), UAS-Crumbs RNAi[330135] (D), UAS-Scribble RNAi[35748] (E) and UAS-Lgl RNAi[109604] (D; yellow or white arrow showing gaps at tricellular or bicellular vertices, respectively).

Scale bars, 20µm (A-F)
