## Supplementary figures and images for "Apical-basal polarity regulates Collagen IV-dependent cell-cell adhesion in the *Drosophila* adipose tissue"

### Supplementary Figure 1

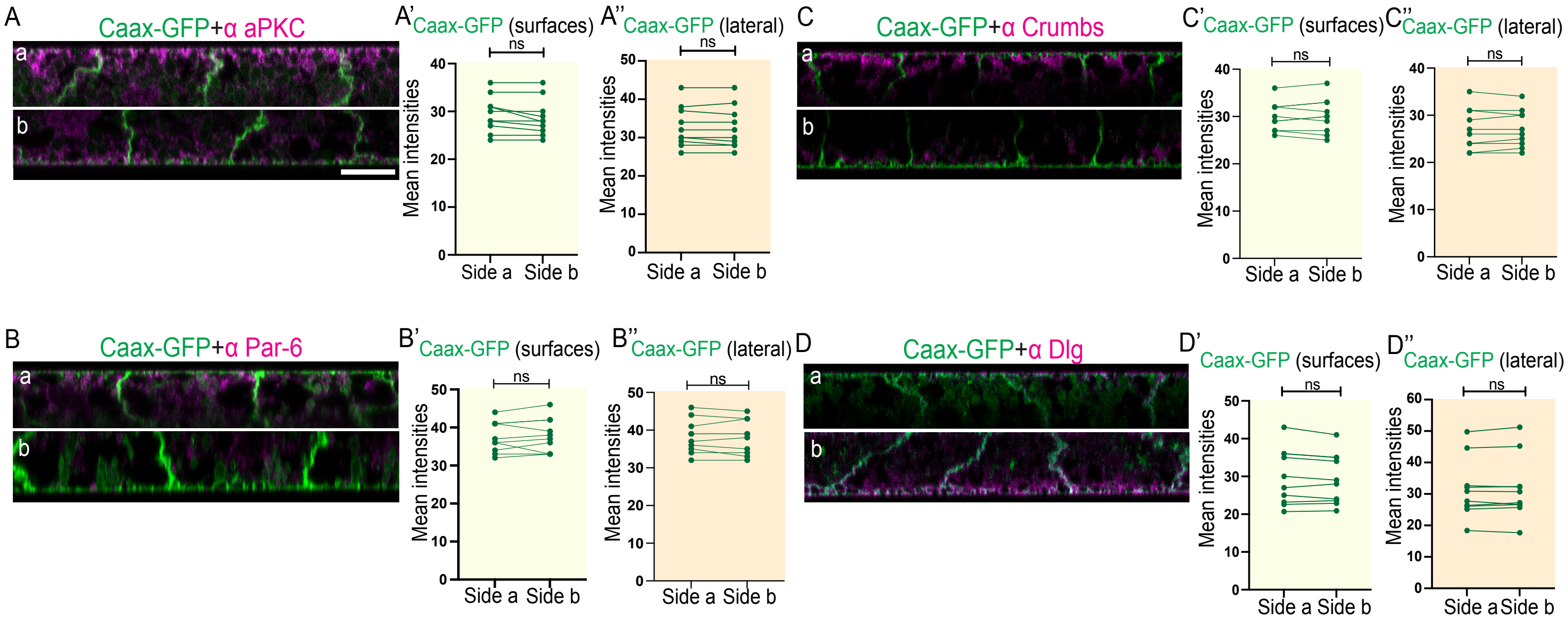

### Supplementary Figure 2

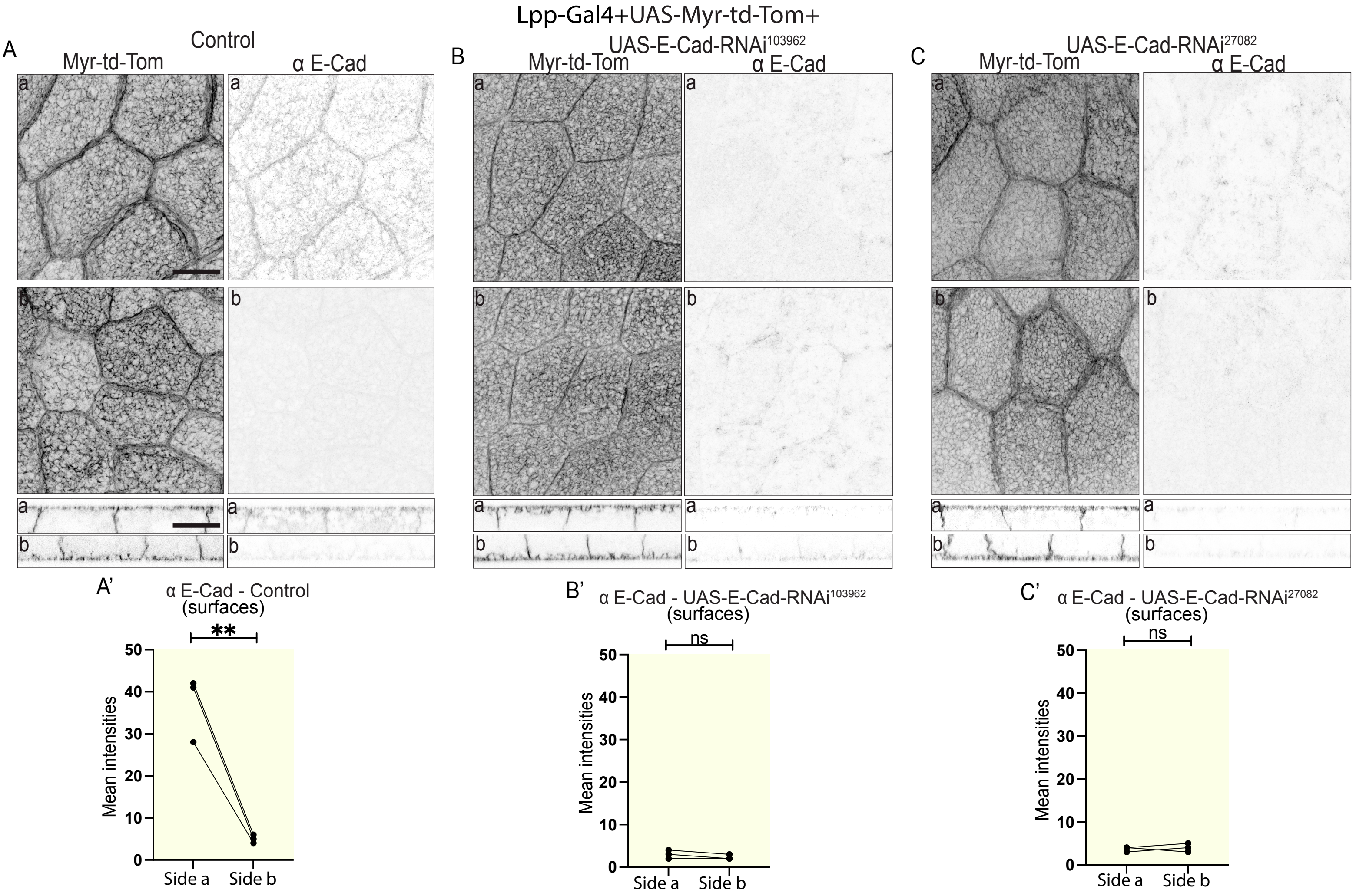

### Supplementary Figure 3

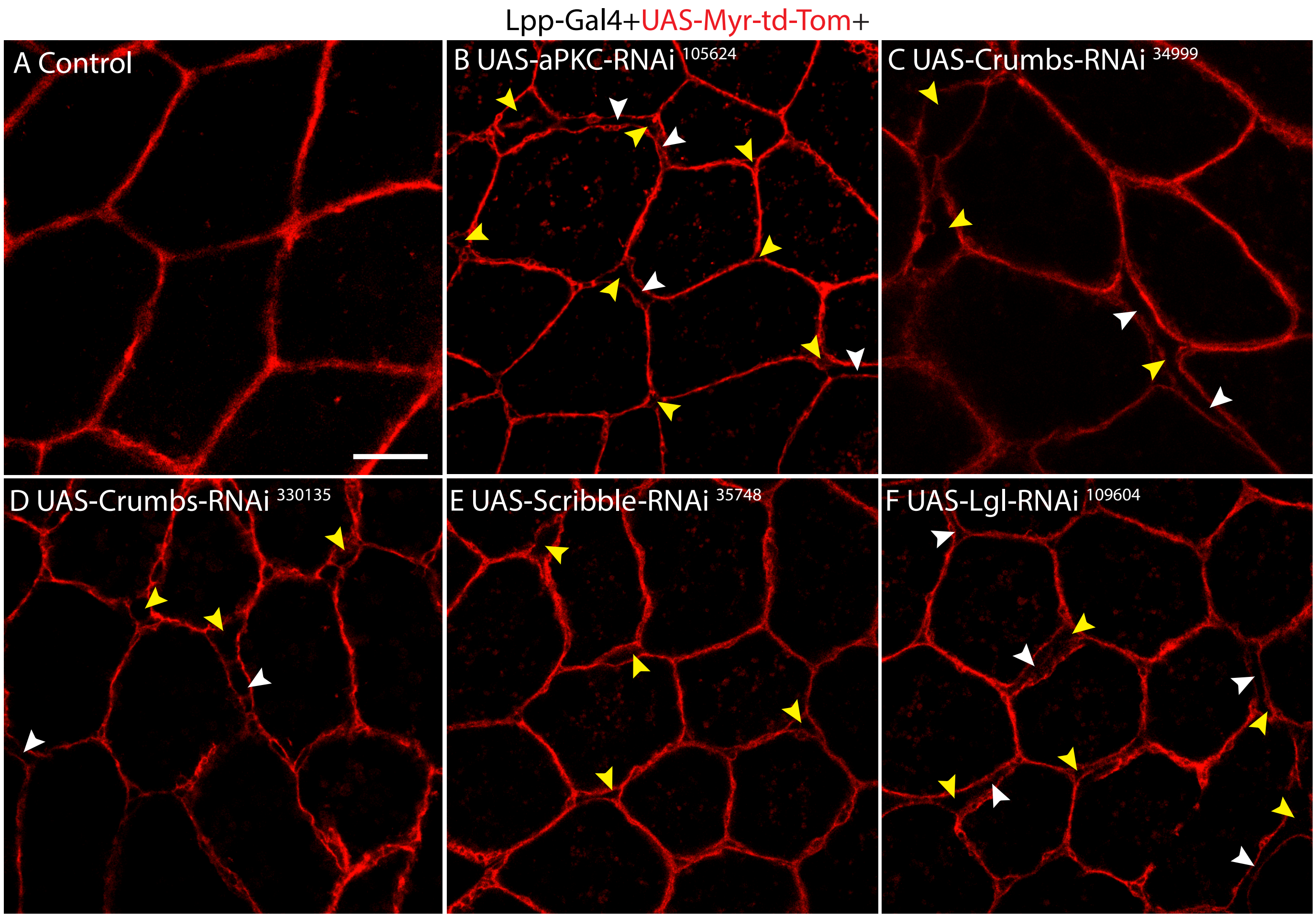
